# Mapping brain function underlying naturalistic motor observation and imitation using high-density diffuse optical tomography

**DOI:** 10.1101/2025.01.21.634109

**Authors:** Dalin Yang, Tessa G. George, Chloe M. Sobolewski, Sophia R. McMorrow, Carolina Pacheco, Kelsey T. King, Rebecca Rochowiak, Evan Daniels-Day, Sung Min Park, Emma Speh, Ari Segel, Deana Crocetti, Alice D. Sperry, Mary Beth Nebel, Bahar Tunçgenç, Rene Vidal, Natasha Marrus, Stewart H. Mostofsky, Adam T. Eggebrecht

**Affiliations:** Washington University School of Medicine, Mallinckrodt Institute of Radiology, St. Louis, Missouri 63110, USA; Johns Hopkins University, Department of Biomedical Engineering, Baltimore, Maryland 21218, USA; Kennedy Krieger Institute, Center for Neurodevelopmental and Imaging Research, Baltimore, Maryland 21205, USA; Nottingham Trent University, Department of Psychology, Nottingham NG1 4FQ, United Kingdom; University of Pennsylvania, Perelman School of Medicine, Department of Radiology, Philadelphia, Pennsylvania 19104, USA; Washington University School of Medicine, Department of Psychiatry, St. Louis, Missouri, 63110, USA; Johns Hopkins University School of Medicine, Department of Neurology, Baltimore, Maryland 21287, USA; Johns Hopkins University School of Medicine, Department of Psychiatry and Behavioral Sciences, Baltimore, Maryland 21287, USA; Washington University School of Engineering, Department of Biomedical Engineering, St. Louis, Missouri 63130, USA; Washington University School of Medicine, Division of Biology and Biomedical Sciences, St. Louis, Missouri 63110, USA; Washington University School of Arts and Science, Department of Physics, St. Louis, Missouri 63130, USA; Washington University School of Engineering, Department of Electrical and System Engineering, St. Louis, Missouri, 63112, USA; Washington University School of Engineering, Department Imaging Sciences Engineering, St. Louis, Missouri, 63112, USA

**Keywords:** Autism spectrum disorder (ASD), non-autistic individuals (NAI), High-density diffuse optical tomography (HD-DOT), Motor observation (OBS), Motor imitation (IM), and Computer-vision-based assessment of motor imitation (CAMI)

## Abstract

**Background:** Autism spectrum disorder (ASD), a condition defined by deficits in social communication, restricted interests, and repetitive behaviors, is associated with early impairments in motor imitation that persist through childhood and into adulthood. Alterations in the mirror neuron system (MNS), crucial for interpreting and imitating actions, may underlie these ASD-associated differences in motor imitation. High-density diffuse optical tomography (HD-DOT) overcomes logistical challenges of functional magnetic resonance imaging to enable identification of neural substrates of naturalistic motor imitation.

**Objective:** We aim to investigate brain function underlying motor observation and imitation in autistic and non-autistic adults. We hypothesize that HD-DOT will reveal greater activation in regions associated with the MNS during motor imitation than motor observation, and that MNS activity will negatively correlate with autistic traits and motor fidelity.

**Methods:** We imaged brain function using HD-DOT in N = 100 participants as they engaged in observing or imitating a sequence of arm movements. Additionally, during imitation, participant movements were simultaneously recorded with 3D cameras for computer-vision-based assessment of motor imitation (CAMI). Cortical responses were estimated using general linear models, and multiple regression was used to test for associations with autistic traits, assessed via the Social Responsiveness Scale-2 (SRS), and imitation fidelity, assessed via CAMI.

**Results:** Both observing and imitating motor movements elicited significant activations in higher-order visual and MNS regions, including the inferior parietal lobule, superior temporal gyrus, and inferior frontal gyrus. Imitation additionally exhibited greater activation in the superior parietal lobule, primary motor cortex, and supplementary motor area. Notably, the right temporal-parietal junction exhibited activation during observation but not during imitation. Higher autistic traits were associated with increased activation during motor observation in the right superior parietal lobule. No significant correlation between brain activation and CAMI scores was observed.

**Conclusions:** Our findings provide robust evidence of shared and task-specific cortical responses underlying motor observation and imitation, emphasizing the differential engagement of MNS regions during motor observation and imitation.

## Introduction

Autism spectrum disorder (ASD) is a neurodevelopmental condition characterized by deficits in social communication and enhanced presentation of restricted interests and repetitive behaviors, with heterogeneous traits across the population^1,2^. Motor impairment is prevalent, with recent estimates suggesting this occurs in up to 88% of autistic children^3–8^. Impairments in motor imitation are particularly notable, as imitation is essential for early prosocial and language development^9,10^. Imitation across a range of communicative behaviors (e.g., vocalizations, arm gestures, facial expressions, and other non-verbal cues) is key components of effective social interaction in neurotypical development^11^. Thus, early imitation deficits detection can hinder social development, and thereby social engagement and communication^12,13^. Additionally, difficulties in visual-motor integration, which enables coordinating visual input with motor execution and is supported in part by the mirror neuron system (MNS) to coordinate visual input with motor execution^14^. Autistic individuals tend to rely more on proprioceptive inputs more than visual cues compared to non-autistic peers when performing movements^15,16^. This increased reliance on proprioception could lead to differences in processing within sensory-motor integration systems, including cortical regions associated with visual processing, motor execution, visual attention, and association regions ^14,17^. Such differences in processing may contribute to the observed motor imitation impairments in autistic individuals ^14^. Therefore, a deeper understanding of the brain function underlying both observation and imitation of natural motor movements is crucial for developing targeted interventions to improve social engagement, communication, and outcomes for autistic and neurotypical individuals.

The current standard in traditional approaches for imitation assessment involved human observation coding (HOC), which introduces bias and requires extensive training^18,19^, thereby making it impractical for use in clinics and home settings. Developing an automatic assessment is valuable but challenging, as human motion is highly heterogeneous and involves spatial and temporal aspects that must quantification capture small changes in imitation to assess these imitation deficits related to autism. Addressing this, Tuncgenc et al. 2021 developed an automatic, Computerized Assessment of Motor Imitation (CAMI) that utilizes metric learning and dynamic time warping for precision measurement of motor imitation fidelity^19^. CAMI quantifies spatiotemporal differences between the movements of a participant and that of a gold standard^20^ and has been validated as a reliable method for assessing motor imitation in school-age children, with CAMI-generated scores being strongly correlated with traditional labor-intensive, subjective HOC based methods^19,21^. Crucially, CAMI methods have proven to be better than HOC at discriminating autistic from non-autistic children, including neurotypical children and those with attention-deficit/hyperactivity disorder (ADHD)^18^. Additionally, CAMI scores have been shown to significantly correlate with measures of autism trait severity (using the Social Responsiveness Scale, Version 2; SRS-2^22^, a well-established clinical metric of autistic traits). Together, these findings demonstrate that CAMI can detect motor imitation difficulties that are specific to autism.

While CAMI has shown promise in detecting diagnostic differences in motor imitation fidelity, the relationships between variability in brain function and motor imitation fidelity remain unknown^23^. Elucidating patterns of brain function underlying motor imitation behaviors may help parse observed heterogeneity in ASD ^12,24^, point to neural mechanisms that lead to deficits, in motor imitation, and guide more individualized treatment planning ^25,26^. The main challenges are due to the physical constraints and limitations of neuroimaging modalities such as functional magnetic resonance imaging (fMRI)^27,28^. Specifically, the MRI environment requires participants to lay supine in a small bore that inherently restricts studies to small-scale movements, such as those of fingers or hands, and it is incompatible to capture brain function during more general, naturalistic motor behaviors ^29^. In contrast, high-density diffuse optical tomography (HD-DOT) provides a naturalistic and open scanning environment that enables the assessment of brain function across the lifespan^30–32^, in school-age autistic children^31,33–36^, or bedside clinical settings such as intensive care units^36^. Importantly, HD-DOT provides signal to noise, anatomical specificity, and spatial resolution approaching that of fMRI^37–39^. Additionally, HD-DOT provides higher spatial resolution ^40–42^ and improved depth sensitivity over a broader cortical field of view than traditional functional near-infrared spectroscopy (fNIRS) methods^37,43^. While previous foundational studies using wearable fNIRS have opened the door to measuring brain function during natural interactions in young children who go on to develop autism^44,45^, more recent advancements in wearable HD-DOT systems^46,47^ highlight the potential for high-fidelity optical neuroimaging to be deployed in both clinic and home settings^48,49^. Given previous success using HD-DOT to map brain function during observation of biological motion ^35^in children with and without autism, herein, we conducted a proof-of-principle study using HD-DOT to detect and localize cortical brain function underlying naturalistic gross motor imitation.

Extant human neuroimaging studies using fMRI suggest motor imitation relies on the MNS, which encompasses the ventral premotor cortex (vPMC), supplementary motor area (SMA), and inferior and superior parietal lobules (IPL, SPL, respectively)^50,51^. These regions, which significantly overlap with praxis networks^52,53^, are crucial for visual-motor integration, i.e., for encoding observed actions and translating them into motor execution. Additionally, the superior temporal sulcus (STS) can be considered part of the MNS, as it plays a role in self-other mapping, facilitating action recognition and intention understanding^50,54–56^. The MNS activates not only when an individual performs an action but also when they observe someone else performing that action^51,57–59^, highlighting the importance of this network in developing abilities to replicate another’s movements and understanding and interpreting the actions of others^50,51,54,56,60–62^. Autism-associated differences in MNS activity have been observed during motor imitation and action observation^26,56,63–69^, with autistic individuals exhibiting reduced MNS activation, impairments in understanding action intentions, and deficits in action prediction ^26,56,63^. Additionally, EEG and fMRI studies found atypical mu suppression, altered neural mirroring, and reduced engagement during social interactions^64–66^. These findings suggest that MNS dysfunction contributes to the social and motor impairments often observed in autistic individuals. Further, the variability in MNS activation has been shown to correlate with autistic traits, suggesting the MNS may underlie key behavioral variability associated with ASD^70^.

Accurate motor imitation also relies on efficient ability to detect and shift attention to others’ actions, which relies on a ventral attention network (VAN) comprising temporoparietal junction (TPJ) and ventral prefrontal cortex^71,72^. It further requires proper application of goal-directed attention and action planning, which relies on a dorsal attention network (DAN) comprising middle temporal V5 area, the intraparietal sulcus (IPS), and frontal eye fields (FEF) regions^73–77^. Difficulties with shifting attention and with goal-directed planning are often reported features of ASD^78,79^; as such, abnormal connectivity of MNS with VAN and DAN may help explain deficits in imitation skills and related behaviors observed in many autistic individuals.

In this study, we used simultaneous HD-DOT and CAMI to investigate how variability in brain function relates to movement observation and imitation fidelity in a sample of autistic and non-autistic adults. Extant literature has largely focused on fine-motor task-based fMRI studies, providing limited insight into the neural activity underlying gross motor observation and imitation in general population, as it could contribute to development of gestures and important for social interaction. Hence, the objective of this study is to address this gap by using limb/truncal movements to assess motor imitation fidelity using CAMI while leveraging HD-DOT to capture the neural correlates of observation and imitation of these movements. Regarding CAMI, we hypothesize that participants in the ASD group will have lower imitation fidelity scores as compared to non-autistic participants. Regarding HD-DOT, we hypothesize that observing and imitating will engage MNS regions including premotor, parietal, and temporal brain regions, with primary motor regions additionally engaged during imitation compared to the resting state. We also hypothesize that regions within the DAN and VAN will be activated both when observing and imitating movements, with stronger brain activation within key regions of the DAN and MNS during motor imitation. Last, we hypothesize that brain response strength within the MNS will be negatively correlated with dimensional measures of autistic traits (i.e., social responsiveness scale, 2nd edition, SRS-2) during observation and imitation, and positively correlated with imitation fidelity (i.e., CAMI scores) during motor imitation.

## Material and Methods

### Participants

We recruited 100 participants (age 30.15 ± 12.97 years, 45 male/55 female, 18 ASD participants, **Table 1**) from the St. Louis, Missouri metropolitan community. The final sample that went into analyses was reduced following exclusions after data quality checks and missing data, as detailed in the Statistical Analyses section below. All participants gave written informed consent to participate in this study, which was approved and carried out in accordance with the Human Research Protection Office at Washington University School of Medicine. All autistic participants had a prior diagnosis from a physician. In the non-autistic individual (NAI) group, the right- to left-handedness ratio was 71:7, while in the ASD cohort, it was 16:3. The population wide ration is 9:1^80^. Additionally, three NAI participants did not report their handedness status.

**Table 1.**
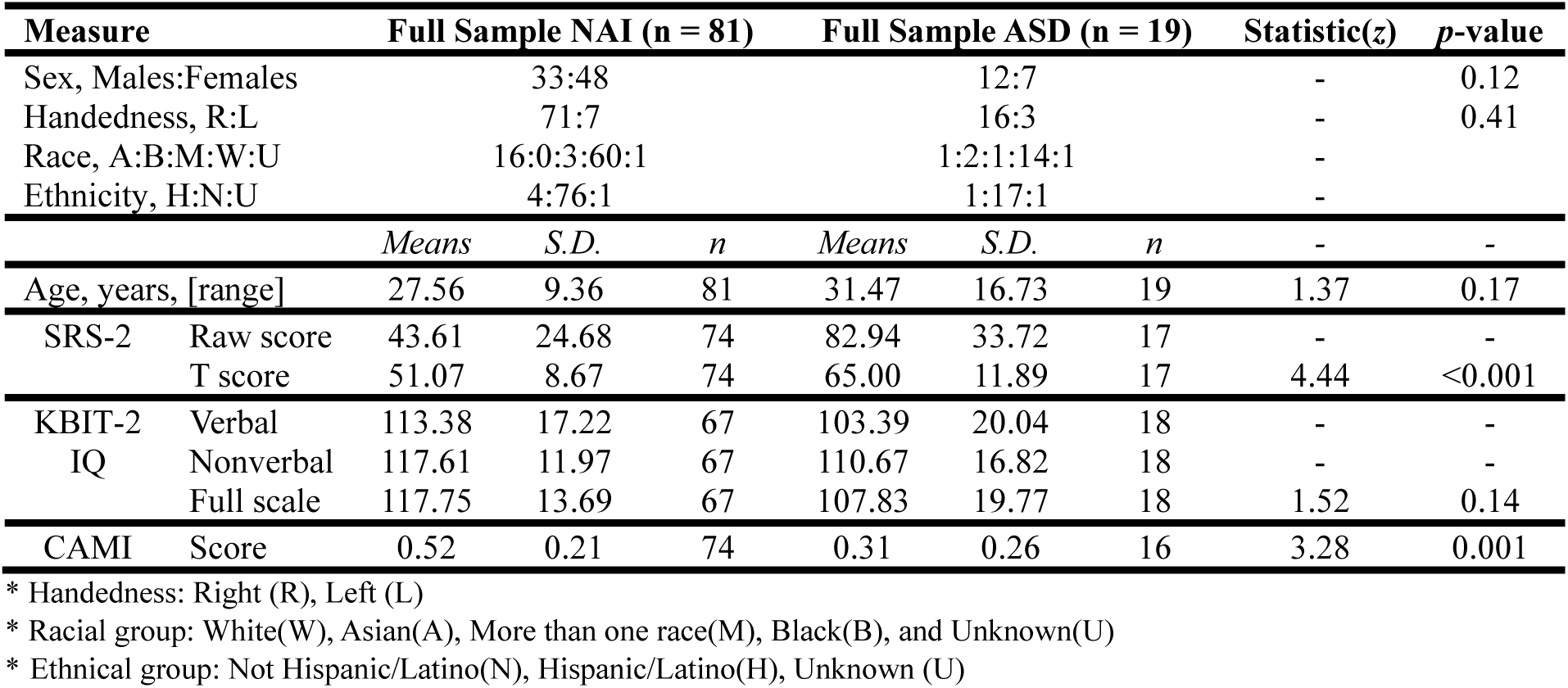
Participant demographics and group characteristic.

### Observation and Imitation Tasks

The motor observation and motor imitation tasks each consisted of six different movement sequences separated by 30-second inter-stimulus-intervals (ISIs; **Figure 1)**. The movement sequences lasted 16-21 seconds and were comprised of 14-18 individual movement types, which were unfamiliar, did not have an end goal, and required moving both arms simultaneously while the participants were seated in a chair. These movements were selected based on prior research indicating that autistic participants experience difficulties performing them ^19^. For the observation task, participants were instructed to passively watch the movement sequences and to remain as still as possible. For the motor imitation task, participants were instructed to mirror the actor’s arm movements as accurately as possible while limiting their head motion (**Figure 1**). Stimuli were presented via the Psychophysics toolbox 3 for MATLAB using custom-built scripts. All movements were performed by the same seated female actor and the stimuli were displayed on a 69 cm monitor placed 110 cm from the nasion in front of the participant. Participant brain function was recorded with HD-DOT during all observation and imitation tasks. Participant imitation fidelity was measured with CAMI using the participants’ motion data recorded during the imitation task. Motion data were recorded using an Xbox Kinect 3D camera located on top of the stimulus presentation monitor and iPi Recorder (iPiSoft4, Moscow, Russia).

**Figure 1:**
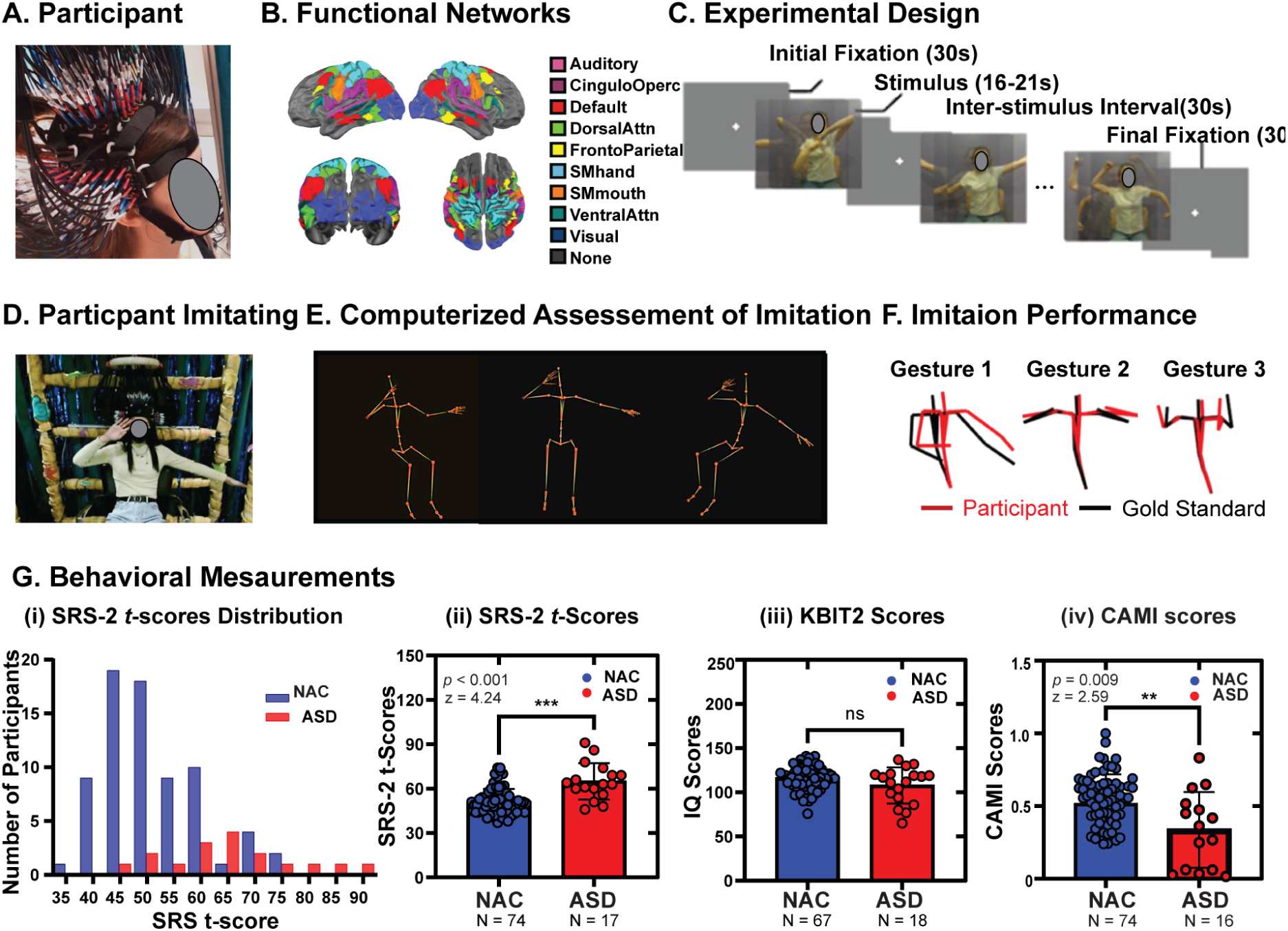
Study Design and Behavioral Assessments. **A** Participant wearing the DOT cap. **B** Field of view from dorsal, posterior, and lateral perspectives and overlapping Gordon parcels colored by functional networks assignment. **C** Experimental design for the motor observation and imitation tasks. **D** Participant performing imitation of movements. **E** Three examples of extracted skeleton of moving participant for CAMI analyses. **F** Comparison of imitation performance with a gold-standard skeleton position during the movements. **G** Behavioral assessments including SRS-2 scores, KBIT-2, and CAMI scores.

### HD-DOT System

The continuous-wave HD-DOT system has been previously described^81^ and was comprised of 128 dual-wavelength optical sources with wavelengths of 685 nm and 830 nm, and 125 avalanche-photodiode detectors. The imaging cap supported multiple source-detector distances (i.e., 11, 24, 33, and 40 mm), providing over 3,500 measurement pairs per wavelength for image reconstruction at a 10 Hz frame rate. The HD-DOT field-of-view covered continuous aspects of temporal, visual, parietal, and prefrontal cortical areas (**Figure 1A, B**). To optimize HD-DOT cap fitting, we utilized anatomical landmarks (e.g., tragus, inion, and nasion) for precise cap alignment. For participants with long hair, the hair on top was parted down the middle and secured into two tight ponytails positioned at the back of the head to minimize interference with the optodes. Participants were then instructed to comb the optodes through their hair until they made even contact with the scalp across the sides and back of the cap. We then secured the cap symmetrically by aligning it with the tragus and fiducials on the cap. Finally, we adjusted the optodes to ensure optimal scalp contact, guided by real-time readouts of data quality metrics such as light levels, noise levels, and the mean light level for each measurement plotted against source-detector separation distance ^43^.

### Behavioral Assessments

The Social Responsiveness Scale, 2^nd^ Edition (SRS-2), provided a measure of quantitative autistic traits, with higher scores corresponding to higher levels of autism traits and lower levels of social reciprocity. SRS scores are continuously distributed throughout the general population. The SRS-2 raw scores were generated based on adult self-report form, and sex-normed SRS T-scores were used in analyses. Participant handedness was determined by the Edinburgh Handedness Inventory (EHI)^82^. Measures of verbal and non-verbal intelligence quotients (IQ) were assessed with the Kaufman Brief Intelligence Test-2 (KBIT-2)^83^. All participants had a composite IQ or verbal IQ > 80.

The CAMI algorithm, described previously^19^, was modified for this study to allow for the imitation task to be used while participants were seated in our fiber-based HD-DOT system. The seated CAMI algorithm considered joints from the upper body as there was no involvement of the lower limbs; all other features of the CAMI analyses remained the same as the previously described methods. The spatial coordinates of 20 joints were extracted from the Xbox Kinect 3D motion sensor depth recordings using iPi Motion Capture Software (iPiSoft 4.6.2, Moscow, Russia). Calculation of CAMI scores consisted of five main steps: (1) preprocessing, (2) automatic joint importance estimation, (3) calculation of the distance feature, (4) calculation of the time features, and (5) calculation of the CAMI score. Preprocessing consisted of (i) defining the hip position as the origin, mapping the sequence of movements to a skeleton of fixed size, (ii) correcting the initial orientation of the participant to match that of the actor (**Figure 1E**), and (iii) the average position of the participants’ joints associated with their spine (Hip, Lower Spine, Middle Spine, and Neck) were corrected to match those of the average position of the gold standard (**Figure 1F**). This alignment step was completed to avoid penalizing participants for their initial posture or sitting position in the cap, as this could vary between participants. The CAMI algorithm yielded an imitation fidelity score ranging between 0 and 1, with higher scores indicating higher fidelity.

### Data pre-processing and Image Reconstruction

HD-DOT data were pre-processed using the NeuroDOT toolbox in MATLAB (https://www.nitrc.org/projects/neurodot). Each of the raw source-detector pair light level measurements were converted to optical density by calculating the log-ratio of the raw measurement data relative to its temporal mean. Measurements with greater than a 7.5% temporal standard deviation in the least noisy (lowest motion) 60s of each run were removed from further analyses. Data were bandpass-filtered between 0.02 and 1.0 Hz using 9-pole Butterworth filters. Then, the superficial signal for each wavelength was estimated as the average across the closest nearest neighbor measurements (11 mm S-D pair separation) and regressed from all measurement channels. The optical density time-traces were then low pass filtered with a cutoff frequency of 0.15 Hz and temporally down sampled from to 1 Hz. As previously described^35^,, a wavelength-dependent sensitivity matrix was computed and inverted to calculate relative changes in absorption at the two wavelengths via reconstruction using a Tikhonov regularization parameter of 0.01 and spatially variant regularization parameter of 0.1. Relative changes in the concentrations of oxygenated, deoxygenated, and total hemoglobin were obtained from the absorption coefficient changes by the spectral decomposition of the extinction coefficients of oxygenated and deoxygenated hemoglobin at the two wavelengths (i.e., 685 and 830 nm). All data were resampled to a 1×1×1 mm^3^ standard MNI atlas using a linear affine transformation for statistical analyses.

### Data Quality Assessment and Motion Censoring

Quantitative data quality assessments included calculating the percentage of good measurements (GM) retained^35^ and the median pulse band signal-to-noise ratio (SNR)^35,43^. As previously described^35^, runs with GM < 80% and/or SNR < 1.0 and were excluded from further analysis to ensure robust data reliability. Global variance in the temporal derivative (GVTD) was used to detect and censor time points with motion-driven artifact in the data^35,39,84^. Briefly, GVTD is calculated as the RMS of the first temporal derivative across the set of first nearest-neighbor measurements for each time point. Higher GVTD values indicate higher global variance in the optical time course and are exquisitely sensitive to motion-based changes in optode-scalp coupling. Motion censoring with GVTD excludes the time points that exceed the GVTD noise threshold. We used a stringent GVTD threshold of 0.1% based on previous studies utilizing GVTD-based motion censoring^35^.

### Statistical Analyses

#### Behavioral and Demographic Measurements

We tested for potential group differences (ASD vs. NAI) in age, SRS-2, KBIT-2, and CAMI scores using Wilcoxon Rank-Sum test, as Shapiro-Wilk tests indicated non-normal distribution. Effect sizes were reported using Rank-Biserial correlation (r), which measures significance based on rank for non-parametric test. Differences in sex and handedness were tested with Fisher’s exact test. To test for group differences in SRS-2 and CAMI scores while controlling for demographic variables, we used analysis of covariance (ANCOVA) to further conduct hypothesis tests to validate the significant differences including age, sex, and handedness as covariates and group index as a fixed factor. Type III Sum of Squares was employed to address potential biases due to unbalanced data and to ensure that covariates were appropriately controlled. Due to the missed handedness (ANCOVA-SRS = 1, ANCOVA-CAMI = 3), there are 90 participants (i.e., 17 ASD and 73 NAI) and 87 participants (i.e., 17 ASD and 73 NAI) used for ANCOVA-SRS and ANCOVA-CAMI tests, respectively. To test the relationship between autistic traits (SRS-2 scores) and imitation fidelity (CAMI scores), we conducted multiple regression analyses for each group. In this study, 83 participants who have both CAMI and SRS scores (i.e., 14 ASD and 69 NAI). Furthermore, we used one-way ANOVA and post-hoc t-tests to examine the impact of race and ethnicity on the data quality (i.e., median SNR and, GM) for all 100 participants across five racial (white *n* = 75, Asian *n* = 17, more than one race *n* = 4, black *n* = 2, and unknown *n* = 2) and three ethnic groups (not Hispanic/Latino *n* = 93, Hispanic/Latino *n* = 5, unknown *n* = 2).

#### Data Inclusion Criteria

After pre-processing and data quality assessment, 78 participants (i.e., 13 ASD and 65 NAI) were identified as having high-quality (i.e., GM > 80%, SNR > 1) observation task data, while 64 participants (i.e., 13 ASD and 51 NAI) had high-quality imitation task data meeting data quality assessment thresholds described in the previous section. Additionally, one participant was excluded due to technical issues with the stimulus presentation. As a result, the final sample consisted of *n* = 77 participants (i.e., 13 ASD and 64 NAI) for motor observation analysis and 63 participants (i.e., 13 ASD and 50 NAI) for motor imitation analysis. Further, nine participants were missing SRS-2 scores and 21 participants were unable to finalize KBIT-2 testing due to non-native English speaking and scheduling difficulties. Therefore, in our final brain-behavior association analyses, *n* =71 participants (i.e., 13 ASD and 58 NAI) have SRS-2 scores and passed data quality thresholds for the observation task, and *n* = 55 participants (i.e., 12 ASD and 43 NAI) have SRS-2 scores and passed data quality threshold for the imitation task. For the CAMI analyses, out of 100 participants, *n* = 89 CAMI scores were valid after exclusion due to damaged video files (*n* = 6), no video recording (*n* = 2), or preprocessing issues with the CAMI data (*n* = 3). Additionally, to avoid possible practice effects from the first imitation run of the motor imitation task, we focus brain-behavior analyses on those 48 participants whose HD-DOT data passed data quality thresholds in the first imitation run.

#### Brain Activation Estimation

We used general linear model (GLM) analyses to estimate the brain response to each task (observation or imitation) using a canonical hemodynamic responses function (HRF) based on HD-DOT data^85^. We present the relative changes in oxygenated hemoglobin (HbO) results as HbO signals exhibit a higher contrast-to-noise ratio compared to deoxygenated hemoglobin (HbR) or total hemoglobin (HbT)^31,84,86^. We used the Gordon cortical parcellation to provide an unbiased spatial sampling for hypothesis-driven analyses. The functionally defined Gordon parcellation provided a framework to couch our analyses within functionally identified parcels of interest. The intersection of each Gordon parcel with the HD-DOT field of view provided 192 parcels associated with nine functional networks (i.e., auditory, frontal-parietal, default mode, dorsal-attention, somatosensory hand, somatosensory mouth, ventral attention, cingulo-opercular, and visual functional networks; **Figure 1B**). Small parcels of size < 500 mm^3^ (approximately equivalent to a 5 mm spherical seed) were excluded from analyses, providing a final parcel count of 118. Based on the Euclidean mean Montreal Neurological Institute (MNI) coordinate of each parcel location and its associated Brodmann area (BA), we selected 58 parcels located in premotor cortex (BA6), inferior parietal lobule (BA39 and 40), inferior frontal gyrus (BA44 and 45), pre-supplementary (BA6) and primary motor area (BA4) to represent the spatially-distributed mirror neuron system (MNS) ^51,54,62^. For the within-task statistics, we used two-tailed paired t-tests to compare the task effect (i.e., observation or imitation) compared to rest (i.e., ISI), with the *p* < 0.05 with a false discovery rate (FDR) correction. We used two-sample t-tests to compare contrast effects between motor observation and motor imitation, with *p* < 0.05 with an FDR correction. Additionally, Cohen’s D is used to measure effect size for the contrasts, including observation vs. rest, imitation vs. rest, and observation vs. imitation. In this study, we first performed brain-wide analyses to test for significant activations during motor observation and imitation. We also investigated which regions were significantly active more during observation vs. imitation. We then highlighted how significantly activated regions are represented within key brain networks previously implicated in observing or imitating motor movements (i.e., MNS, DAN, VAN). Finally, we investigated if cortical activations during observation were associated with SRS scores and if cortical activations during imitation were associated with either SRS scores or simultaneously measured imitation fidelity (CAMI scores).

#### Brain Behavioral Association Analysis

We calculated Pearson correlations between the parcel-specific brain responses to motor observation and motor imitation with the SRS-2 *t*-scores to investigate brain-behavior correlations relative to autistic traits. We also calculated the Pearson correlation between the motor imitation responses and the CAMI scores to assess brain-behavior relationships relative to motor imitation fidelity. Further, we used multiple regression analysis to examine how demographic factors (i.e., age, sex, and handedness) might affect the correlation between above brain and behavioral associations. More specifically, we assessed the effects of sex, age, and handedness, along with behavioral measurements (i.e., SRS-2 or CAMI scores), using brain activation data that passed the *p* < 0.05 threshold (with and without FDR correction) as the dependent variables. Overall, Pearson correlation is helpful for understanding fundamental relationships between brain and autistic traits (SRS-2 *t*-scores and CAMI scores), while multiple regression provides a more comprehensive perspective, accounting for all potential effects of multiple demographic predictors (i.e., age, sex, and handedness) and autistic traits SRS-2 *t*-scores and CAMI scores).

## Results

### Behavioral Measures

The participant demographics and group characteristics are listed in Table 1. In our samples, autistic participants exhibited significantly higher scores on the SRS-2 compared to non-autistic individuals (NAI; **Figure 1G i, ii**; NAI *n* = 74, ASD *n* = 17, *z* = 4.24, *r* = 0.91, *p* < 0.001). There were no significant differences between the NAI and ASD groups in verbal or non-verbal KBIT scores (**Figure 1G iii**, NAI *n* = 67, ASD *n* = 18, *z* = 1.29, *r* = 3.98, *p* = 0.20). In line with previous CAMI studies and our hypothesis, the NAI group exhibited significantly higher CAMI scores compared to the ASD participants (**Figure 1G iv**; NAI *n* = 74, ASD *n* = 16, *z* = 2.59, *r* = 5.41*, p* = 0.009), indicating that the NAI group had generally higher motor imitation fidelity scores than the ASD group. Similarly, our ANCOVA results showed that a significant group difference remains in SRS scores (F = 25.76, *p* < 0.001) and CAMI scores (F = 5.53, *p* = 0.02) even after controlling for demographic variables, including age, sex, and handedness. Interestingly, we observed a significant effect of sex (F = 4.33, *p* = 0.04) on the CAMI scores across the entire sample with males scoring higher than females (F = 4.33, *p* = 0.04). No significant group and sex interaction (F = 0.04, *p* = 0.85) was found, indicating that the influence of sex on CAMI scores was consistent across both groups.

Regarding handedness, the ratio of right- to left-handed individuals in our overall sample was 10.3% (10/97), closely aligning with the general population’s handedness distribution of 10.6%^80^. Moreover, there was no significant difference between the NAI and ASD groups with respect to age (ASD *n* = 19, NAI *n* = 81, *t* = 1.37, *p* = 0.17), sex (ASD *n* = 19, NAI *n* = 81, *p* = 0.12), or handedness (ASD *n* = 19, NAI *n* = 78, *p* = 0.41). Furthermore, significant differences in GM were found across five race groups (F = 10.01, *p* < 0.001, white *n* = 75, Asian *n* = 17, more than one race *n* = 4, black *n* = 2, and unknown *n* = 2), with post-hoc t-tests revealing a significant difference between Asian and white groups (Cohen’s D = -1.21, *p* < 0.001), while a significant difference in median SNR was found across races (F = 2.86, *p* = 0.04). However, no significant differences were observed in either median SNR (F = 0.68, *p* = 0.51) or GM (F = 0.61, *p* = 0.55) across ethnic groups (not Hispanic/Latino *n* = 93, Hispanic/Latino *n* = 5, unknown *n* = 2), and although there was an overall difference in Med SNR across racial groups, no pairwise comparisons were significant in the post-hoc analysis.

### Neural Correlates of Motor Observation and Imitation

During motor observation, significant activation (**Figure 2**, FDR corrected *p* < 0.05) was found in visual, temporal, and prefrontal cortex compared to the resting state. Bilateral regions throughout visual cortex (*t* > 5.74, Cohen’s D > 0.66, FDR corrected *p* < 0.001), including the occipital lobe, were highly engaged during observation task. In temporal cortex, the superior temporal gyrus (STG, *t* = 6.05, Cohen’s D = 0.69, FDR corrected *p* = 1 × 10^-6^), medial temporal gyrus (MTG, *t* = 5.53, Cohen’s D = 0.63, FDR corrected *p* = 0.006) and fusiform gyrus (FFG, *t* = 5.10, Cohen’s D = 0.58, FDR corrected *p* = 0.003) were also strongly recruited. Bilateral inferior frontal gyrus (IFG, *t* = 4.77, Cohen’s D = 0.58, FDR corrected *p* = 0.003) and inferior parietal lobe (IPL, *t* = 4.62, Cohen’s D = 0.53, FDR corrected *p* = 0.002) were also strongly activated during observation compared to the resting state.

**Figure 2:**
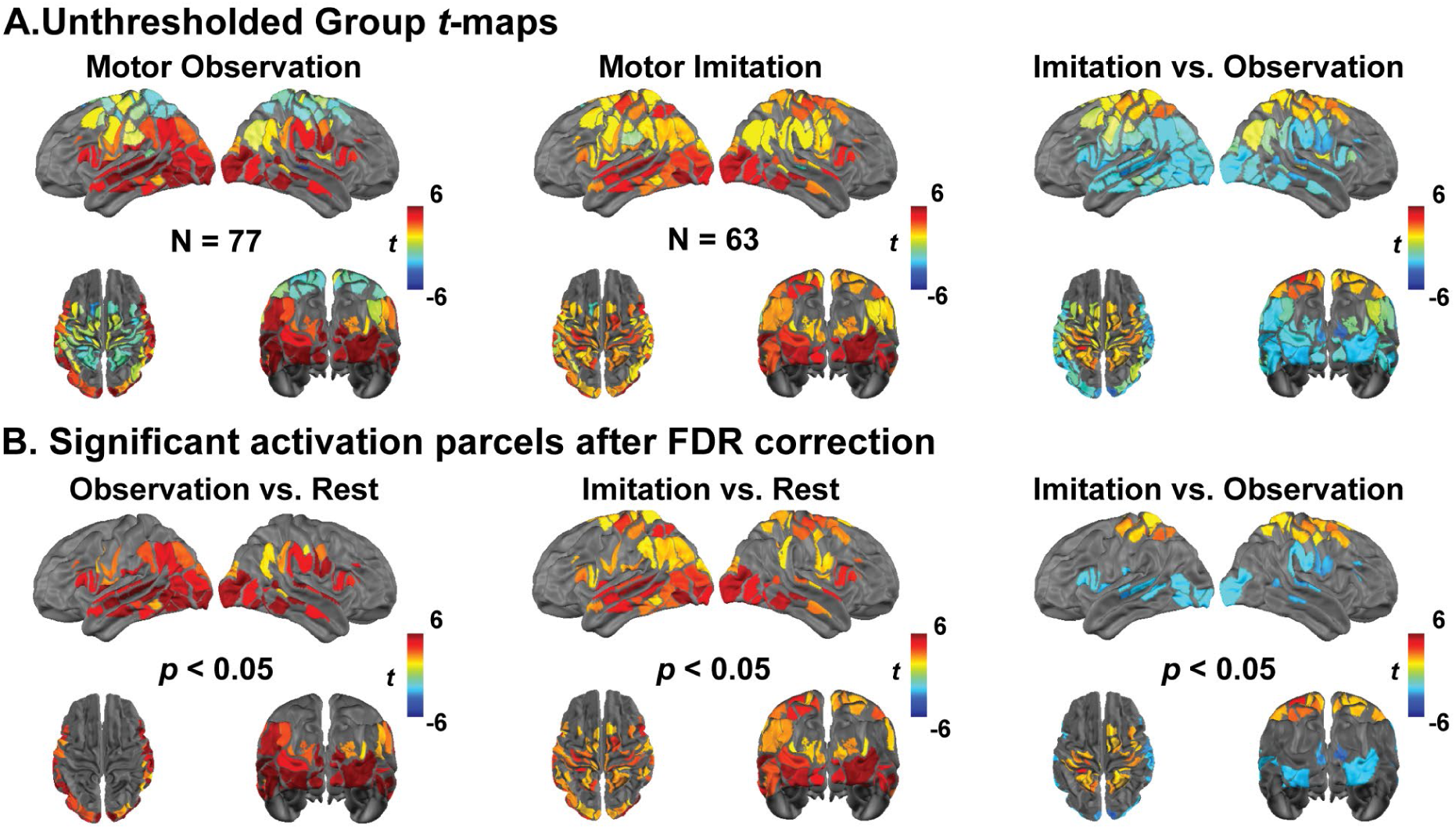
Neural correlates of observation and imitation. **A.** Group-level t-maps for motor observation (*n* = 77), motor imitation (*n* = 63), and their contrast (imitation vs. observation). **B.** FDR-corrected t-maps highlighting statistically significant regions with false discovery rate correction (FDR corrected *p* < 0.05) applied to account for multiple comparisons.

During motor imitation, additional regions of activation extended beyond above regions (i.e., STG, MTG, FFG, and IFG) and included key motor-related areas (**Figure 2**, FDR corrected *p* < 0.05). The primary motor cortex (M1, *t* = 3.18, Cohen’s D = 0.40, FDR corrected *p* = 0.009), premotor cortex (PMC, *t* = 3.19, Cohen’s D = 0.40, FDR corrected *p* = 0.02), and supplementary motor area (SMA, *t* = 2.91, Cohen’s D = 0.37, FDR corrected *p* = 0.02) showed significant engagement during motor imitation task. Additionally, the frontal eye field (FEF, *t* = 2.93, Cohen’s D = 0.37, FDR corrected *p* = 0.01) and superior parietal lobules (SPL, *t* = 3.39, Cohen’s D = 0.43, FDR corrected *p* = 0.01) exhibited significant activations compared to the resting state.

Conjunction analyses of overlapping activation for imitation and observation task states revealed activation in MNS regions, including IPL, STG and IFG, as well as in bilateral occipital lobe, FFG and MTG regions (**Figure 3**). Despite these similarities, consistent with above, direct contrast of imitation vs. observation task states revealed significant activation (**Figure 2 B**, FDR corrected *p* < 0.05) in motor/premotor regions, including bilateral M1 (*t* = 2.91, Cohen’s D = 0.49, FDR corrected *p* = 0.02), PMC (*t* = 2.50, Cohen’s D = 0.42, FDR corrected *p* = 0.03), and SMA (*t* = 2.43, Cohen’s D = 0.41, FDR corrected *p* = 0.03; **Figure 4**, imitation > observation). In contrast, direct contrast of motor observation vs. imitation exhibited significant activation (**Figure 2B**, FDR corrected *p* < 0.05) principally in posterior cortical regions: visual and temporal cortex, including bilateral visual cortex (*t* = 2.48, Cohen’s D = 0.41, FDR corrected *p* = 0.03), STG (*t* = 2.73, Cohen’s D = 0.45, FDR corrected *p* = 0.02) and MTG (*t* = 2.48, Cohen’s D = 0.40, FDR corrected *p* = 0.03); additionally, IFG activation was also observed for observation vs. imitation (*t* = 2.31, Cohen’s D = 0.38, FDR corrected *p* = 0.04; **Figure 5**, observation > imitation). These results revealed that motor imitation strongly activated motor-related regions, including M1, SMA, PMC, and parietal areas (**Supplementary Figure 1**), which remained inactive during the observation task compared to the resting state. In contrast, observation primarily engages action understanding and perceptual processing brain regions (**Supplementary Figure 2**), with greater activation in the left IFG and right temporal parietal junction (TPJ), which remain inactive during the imitation task. This highlights the brain regions for analyzing rather than executing movement in the context of motor imitation. Notably, compared to observing, imitating led to significantly greater activation in extended MNS areas, including M1 and SMA (**Supplementary Figure 3B, C**). To further investigate the observed brain activations from a functional network perspective, we quantified the number of cortical parcels activated within each network for each task (**Supplementary Figure 4**). Most brain networks were dynamically engaged during motor observation and imitation tasks. The dorsal attention network and the somatosensory hand network (*t* = 2.63, Cohen’s D = 0.45, FDR corrected *p* = 0.02) exhibited larger involvement during imitation than observation (**Supplementary Figure 4D, Supplementary Figure 5**). In contrast, the visual (*t* = 1.86, Cohen’s D = 0.31, FDR corrected *p* = 0.04) and ventral attention network (VAN, *t* = 2.39, Cohen’s D = 0.39, FDR corrected *p* = 0.03) exhibited greater recruitment during observation than imitation (**Supplementary Figure 4D, Supplementary Figure 5**). Overall, these results support our hypothesis that both observation and imitation engage the MNS, with imitation showing stronger activation in extended MNS regions, including M1 and SMA. Additionally, as predicted, DAN showed greater activation during imitation, while VAN was more engaged during observation, confirming distinct network dynamics between the two tasks.

**Figure 3:**
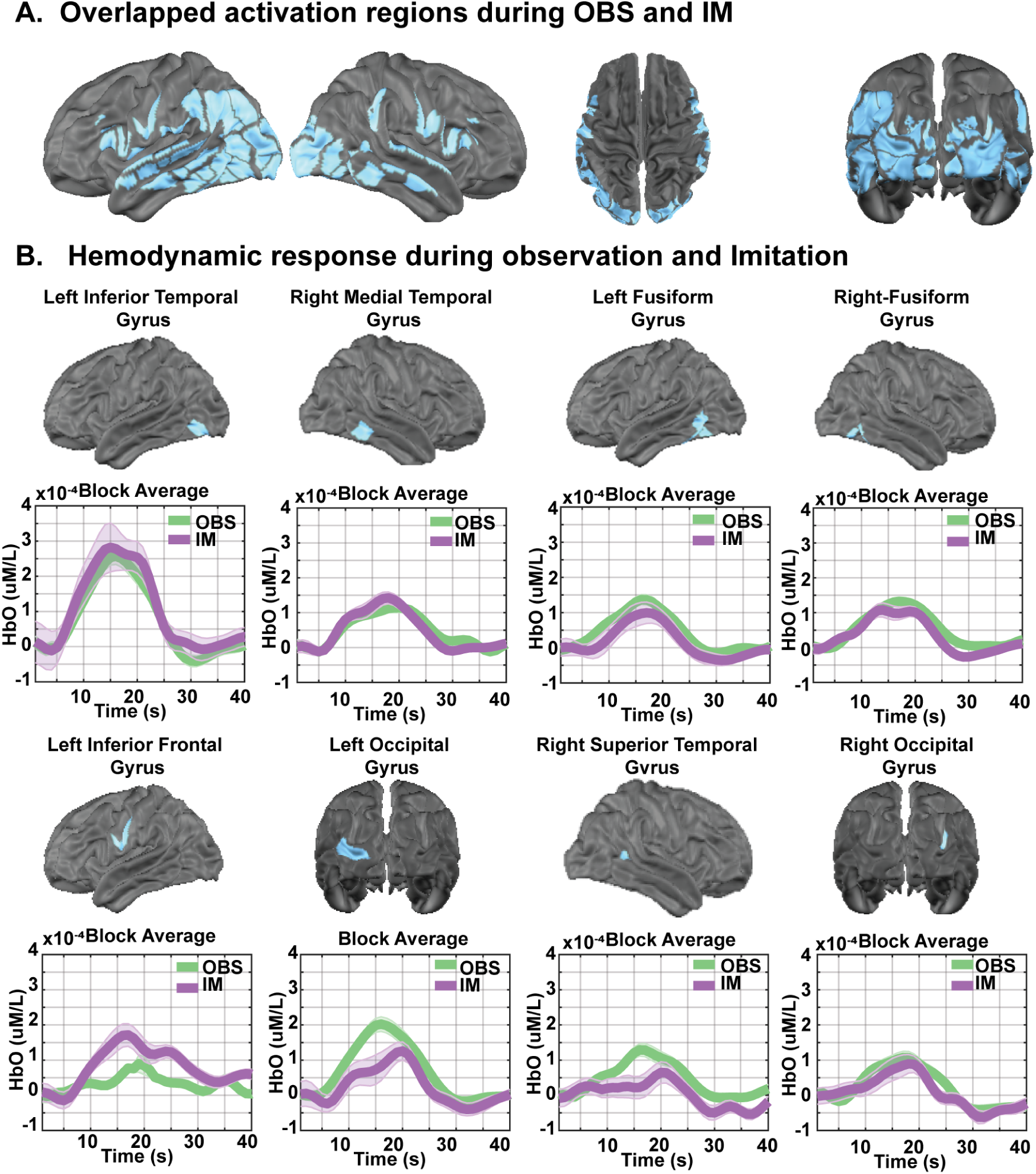
Brain regions exhibiting significant responses to both motor observation and imitation. **A.** Brain regions with overlapping significant activation (FDR corrected *p* < 0.05) identified through conjunction analysis. **B.** Temporal profiles of the HbO (oxygenated hemoglobin) hemodynamic response for each task, with shaded areas indicating the standard error of the mean across participants.

**Figure 4:**
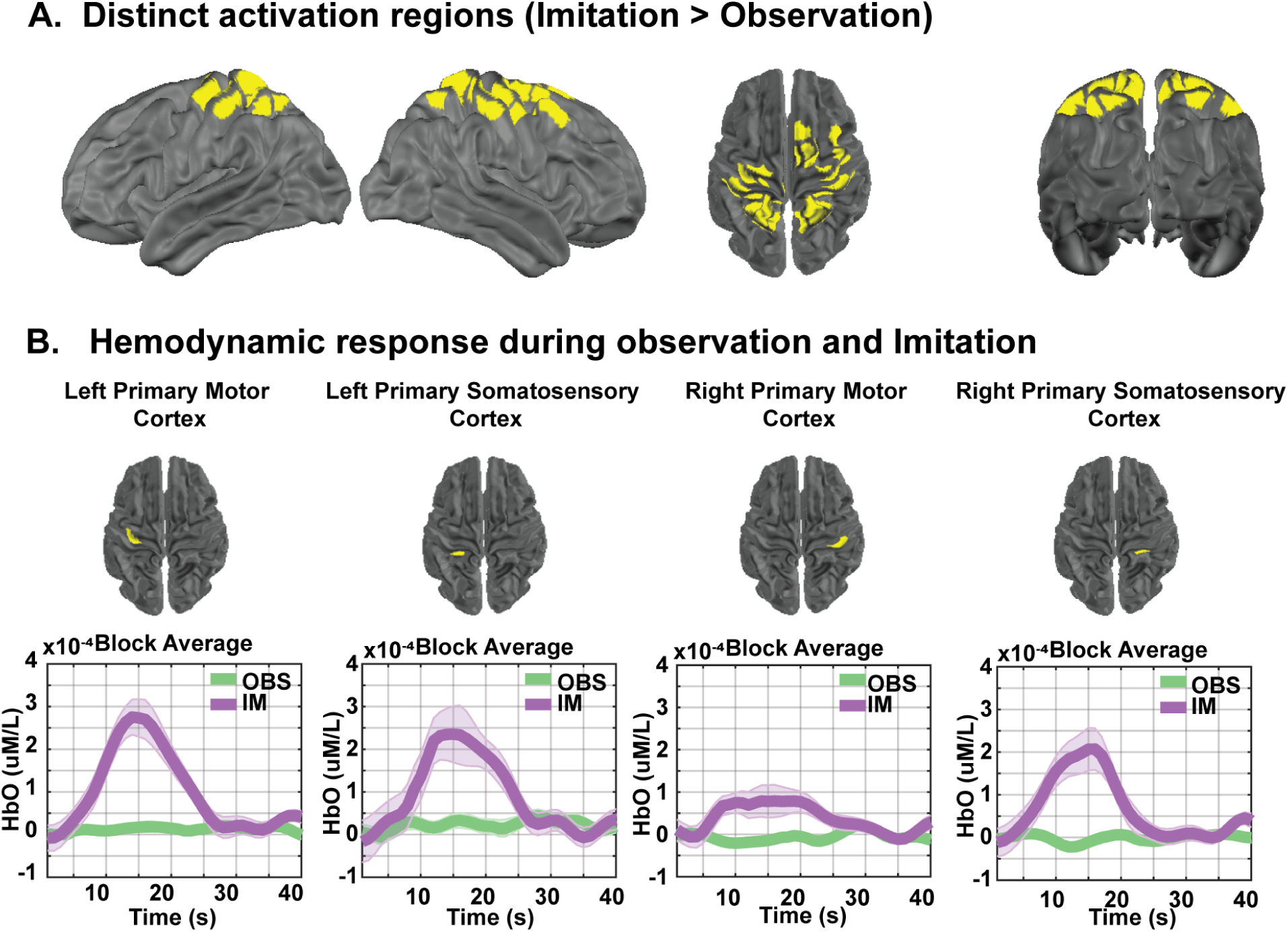
Brain regions exhibiting significantly greater responses during motor imitation than motor observation (FDR corrected *p* < 0.05). **A.** Brain regions exhibit significantly greater activation during motor imitation compared to motor observation task. **B.** Temporal profiles of the HbO (oxygenated hemoglobin) hemodynamic response for each task, with shaded areas representing the standard error of the mean across participants.

**Figure 5:**
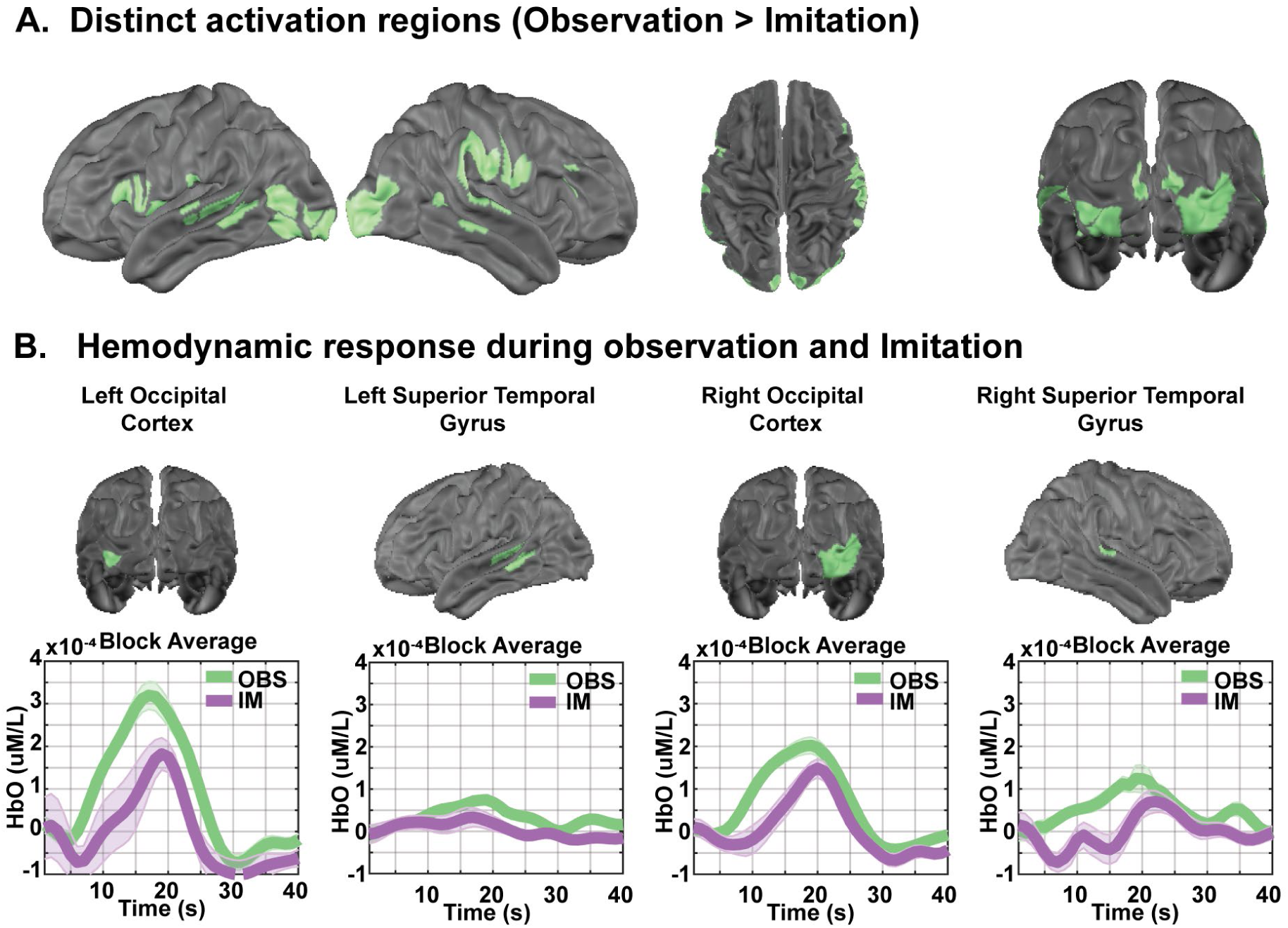
Brain regions exhibiting significantly greater responses during motor observation than motor imitation (FDR corrected *p* < 0.05). **A.** Brain regions exhibit significantly greater activation during motor observation compared to motor imitation. **B.** Temporal profiles of the HbO (oxygenated hemoglobin) hemodynamic response for each task, with shaded areas representing the standard error of the mean across participants.

### Associations with SRS-2

The Pearson correlation analysis revealed a significant positive correlation between SRS-2 t-scores and brain activation in the right SPL during the observation task (**Figure 6A**, *r* = 0.46, *p* = 3.4 × 10^-5^, FDR-corrected *p* = 0.007), such that individuals with higher autistic traits exhibited greater activation in the r-SPL during motor observation. Our overall multiple regression model, including SRS and demographic variables (i.e., age, sex, and handedness), predicted the brain region of r-SPL (F (70) = 5.18, R^2^ = 0.24, adjusted R^2^ = 0.19, *p* < 0.001). Additionally, among these predictors, SRS-2 scores significantly predicted (i.e., *β* = 0.43, *t* = 3.94, *p* < 0.001) brain activation in right SPL, while demographic variables did not (i.e., age: *β* = -0.10, *t* = 0.94, *p* = 0.34; sex: *β* = 0.15, *t* = 1.36, *p* < 0.18), and handedness: *β* = -0.03, *t* = 0.25, *p* = 0.80). We found additional regional associations that were sub-threshold for significance and are presenting them as hypothesis-generating results for future studies that may be more strongly powered to test these effects. Specifically, we observed strong (but not significant enough to pass FDR correction) positive correlations between SRS-2 t-scores and brain activation in the left IPL (*r* = 0.24, *p* = 0.045), bilateral SMA (left SMA: *r* = 0.25, *p* = 0.03; right SMA: *r* = 0.28, *p* = 0.01) during the observation task. Moreover, the Pearson correlation analysis has shown that activation magnitudes in the bilateral occipital lobe (left occipital lobe: *r* = -0.31, *p* = 0.009; right occipital lobe: *r* = -0.24, *p* = 0.04) exhibited a negative correlation with SRS-2 scores (**Figure 6A**). Regarding the multiple regression analysis, the overall model significantly predicted the brain activation in the right IPL (F (70) = 5.18, R^2^ = 0.24, adjusted R^2^ = 0.19, *p* < 0.001) and right occipital lobe (F (70) = 6.42, R^2^ = 0.28, adjusted R^2^ = 0.24, *p* < 0.001). Additionally, SRS-2 scores are the primary predictor for the brain activation prediction (right IPL: *β* = 0.433, *t* = 3.94, *p* < 0.001; right occipital lobe: *β* = -0.40, *t* = 3.75, *p* < 0.001) while controlling for demographic variables (i.e., age, sex, and handedness). However, after including demographic variables, the overall model did not reach statistical significance in the other brain regions, such as the bilateral SMA (right SMA: F (70) = 1.72, R^2^ = 0.09, adjusted R^2^ = 0.04, *p* = 0.16; left SMA: F (70) = 2.315, R^2^ = 0.12, adjusted R^2^ = 0.07, *p* = 0.07), left IPL (F(70) = 1.06, R^2^ = 0.06, adjusted R^2^ = 0.003, *p* = 0.39), and left occipital lobe (F(70) = 2.09, R^2^ = 0.11, adjusted R^2^ = 0.06, *p* = 0.09). Nonetheless, SRS still demonstrated significant predictive power in the right SMA (*β* = 0.29, *t* = 2.40, *p* = 0.02) and left occipital lobe (*β* = -0.32, *t* = 2.66, *p* = 0.01). Additionally, we found a significant sex effect contribution (*β* = -0.28, *t* = 2.60, *p* = 0.01) to prediction of brain activation in right occipital lobe along with SRS-2 scores.

**Figure 6.**
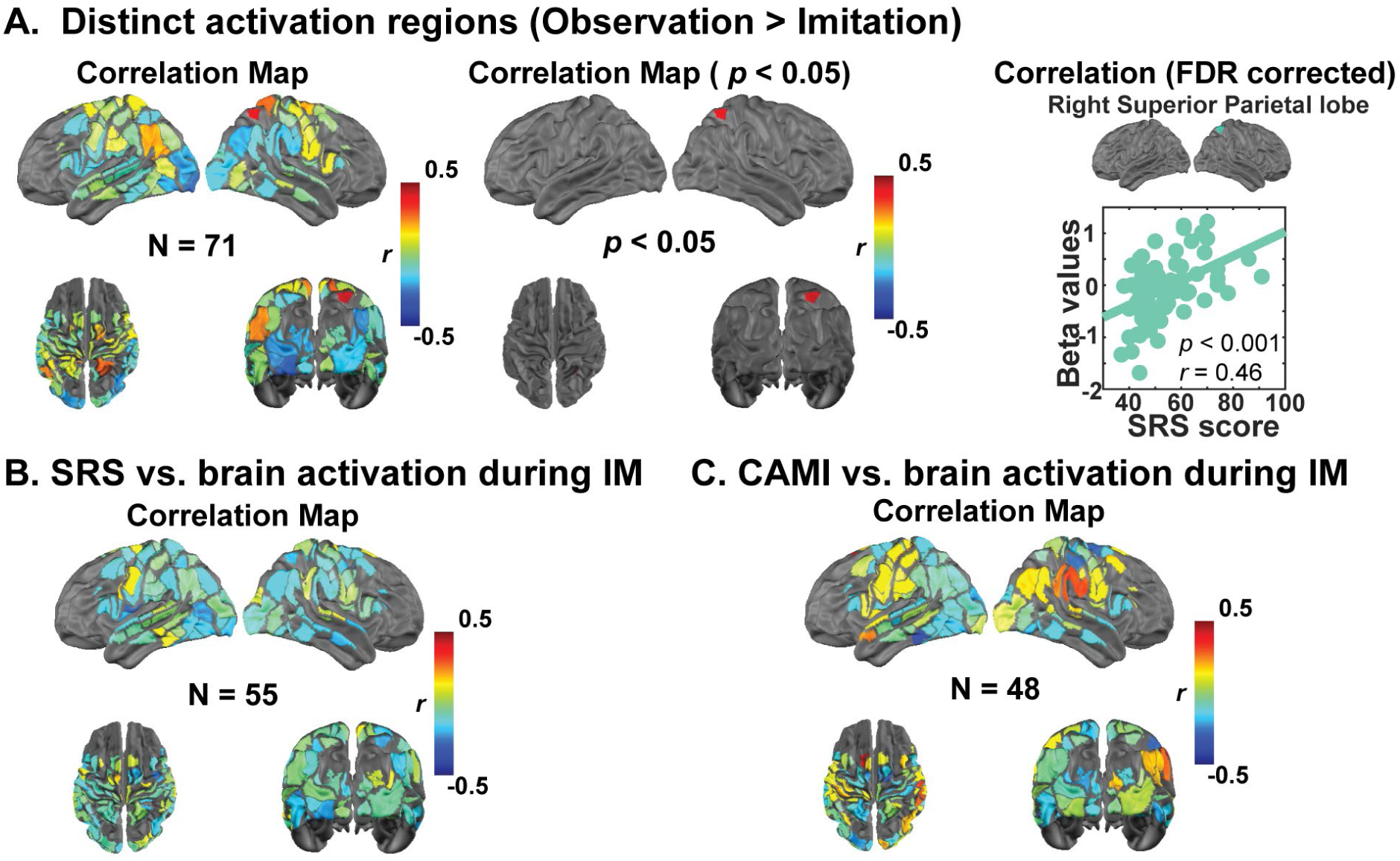
Brain-behavioral association between SRS-2 t-scores and brain activation. **A (i)** Correlation map of brain activation during motor observation with SRS-2 in 71 participants**, (ii)** the brain regions which correlation *p* < 0.05, and **(iii)** the correlation passed the FDR correction (*r* = 0.472, and FDR corrected *p* = 0.007, and)**. B** Correlation map of brain activation during motor imitation with SRS-2 in 55 participants. **C** Brain-behavioral association between CAMI and brain activation during motor imitation task for 48 participants.

The Pearson correlation analysis revealed a negative correlation between SRS-2 t-scores and brain activation in the left subcentral gyrus (**Figure 6B**, *r* = -0.30, *p* = 0.03) during the imitation task. The multiple regression results indicate that the overall model significantly explains variance in the left subcentral gyrus (F (53) = 3.33, R² = 0.21, adjusted R² = 0.15, *p* = 0.02). Similarly, SRS remains the significant predictor (*β* = -0.38, *t* = 2.857, *p* = 0.006), and sex also shows a significant prediction power (*β* = -0.36, *t* = 2.68, *p* < 0.01), whereas age (*β* = 0.10, *t* = 0.81, *p* = 0.43) and handedness (*β* = -0.10, *t* = 0.77, *p* = 0.45) did not contribute significantly to the model. These findings suggest that individuals with higher autistic traits may show diminished brain responses in left subcentral gyrus during motor imitation tasks. However, these correlations did not survive FDR correction.

Our hypothesis-driven analyses focusing on the mirror neuron system (MNS) revealed negative correlations between SRS-2 t-scores and activation in MNS-related regions, though these did not survive FDR correction. This suggests that individuals with elevated autistic traits may exhibit lower activity in these areas during social tasks. No significant correlations were found between SRS-2 scores and activation in the dorsal or ventral attention networks (DAN or VAN) during imitation.

### Associations with CAMI

Our results indicate that no brain activity correlations with CAMI scores remained statistically significant after applying FDR correction (**Figure 6C**). We are presenting sub-FDR-threshold results as hypothesis-generating information for future studies. First, positive correlations were observed in left FEF (*r* = 0.43, *p* = 0.002) and right IPL (*r* = 0.31, *p* = 0.03, within MNS). Conversely, negative correlations were identified in right M1 (*r* = -0.29, *p* = 0.04, within MNS) and left FFG (*r* = 0.34, *p* = 0.02). Separate multiple regression models were conducted per brain region of interest with CAMI scores as the predictor, and age, sex, and handedness as the control variables. The overall model significantly predicted the brain activation in the region in left FEF (F(46) = 4.96, R^2^ = 0.32, adjusted R^2^ = 0.26, *p* = 0.002), right IPL (F(46) = 2.83, R^2^ = 0.21, adjusted R^2^ = 0.14, *p* = 0.04).Among these predictors, the CAMI scores significantly predicted the brain variance in left FEF (*β* = 0.439, *t* = 3.30, *p* = 0.002), right IPL (*β* = 0.37, *t* = 2.58, *p* = 0.01). Similarly, the CAMI score independently contributes to predicting activation in right M1 (*β* = -0.31, *t* = 2.10, *p* = 0.04) and left FFG (*β* = -0.31, *t* = 2.13, *p* = 0.04). The whole model with demographic variables did not explain the brain activation significantly in right M1 (F (46) = 2.28, R^2^ = 0.18, adjusted R^2^ = 0.10, *p* = 0.13), and left FFG (F (46) = 1.90, R^2^=0.21, adjusted R^2^ = 0.14, *p* = 0.07). Interestingly, we also found age significantly contributes to explain the variance of brain activation in the left FEF along with the primary factor (i.e., CAMI score; *β* = -0.39, *t* = 2.97, *p* = 0.005). Additionally, handedness significantly predicted the correlation of brain activity in the right IPL with CAMI scores (*β* = -0.32, *t* = 2.19, *p* = 0.03).

## Discussion

This study investigated brain activation underlying naturalistic motor observation and imitation under more naturalistic conditions than possible using fMRI. Our findings revealing both the shared and task-specific brain activation patterns for observation and imitation tasks. Consistent with our hypothesis, we found that the core MNS regions, including the superior temporal gyrus (STG), inferior parietal lobule (IPL), and inferior frontal gyrus (IFG), were engaged during both tasks, with imitation evoking significantly greater activation in the primary motor cortex (M1), premotor cortex (PMC), and supplementary motor area (SMA), underscoring their involvements in coordinating and executing motor imitative actions. Additionally, as we hypothesized, regions within the dorsal (DAN) and ventral (VAN) attention networks were engaged during both tasks, with stronger activation in key DAN and MNS regions during motor imitation. In contrast, the temporoparietal junction (TPJ) was engaged during observation but remained less active during imitation. Contrary to our hypothesis, higher autistic traits (SRS-2 scores) were associated with increased activation in the right SPL during observation and decreased activation in the left subcentral gyrus during imitation, and CAMI scores showed only trend-level associations with brain activity. These findings provide valuable insights into how brain activation variability in specific brain regions may contribute to motor imitation deficits and their association with autistic traits.

During the motor observation task (**Figure 2, 5, and Supplementary Figure 2**), significant activations were observed in the STG, IPL, IFG, and TPJ. These regions are closely associated with the mirror neuron system (MNS) and ventral attention network (VAN) and are known to facilitate the processing of observed actions^51,57,91^. The STG, particularly the posterior STG, exhibited strong activity, which aligns with its role in mapping observed movements actions onto previously neural spatial-temporal representations, as it is usually considered as the visual input of MNS^51,57,66,92,93^. The right TPJ, part of the VAN, was strongly activated during the motor observation task but not during the motor imitation task. This aligns with its role in stimulus-driven attention and social perception, particularly when the participant observes the actor’s movements without the task demand of imitating^94^. Additionally, the right TPJ is crucial for understanding others’ actions, intentions, and goals, making it highly relevant for observational learning^94,95^. However, during imitation, its activity diminishes as the cognitive focus shifts from action interpretation to direct motor execution. This transition from motor observation to first-person motor mapping is supported by the involvement of the IFG and IPL, key regions of the MNS, which facilitate action understanding and execution ^96,97^. The IFG, particularly, has been shown to play a role in action imitation and motor planning, while the IPL contributes to sensorimotor integration with spatial-temporal representations of motor intention^98^. Together, these areas shift the neural representation from a passive observational stance to active, embodied execution during imitation ^91,96,97^.

Motor imitation elicited significantly stronger activations in primary motor cortex (M1), supplementary motor area (SMA), dorsal premotor cortex (PMC), and the intraparietal sulcus (IPS) compared to motor observation (**Figure 2**, **4, and Supplementary 1**). The SMA and PMC play key roles in motor planning and movement coordination ^99,100^, explaining their enhanced activity when participants transitioned from passively observing movements to actively reproducing them. Additionally, the posterior parietal cortex, particularly the IPS, plays a crucial role in visual-motor integration by combining of visual and spatial information to necessary to planning and executing goal-directed movements ^101^. A direct contrast between imitation and observation revealed additional recruitment from posterior temporal-parietal regions (STG, IPL) and frontal-motor regions (M1, SMA, PMC, IFG). This additional neural recruitment is expected, as imitation requires additional motor control mechanisms beyond those used for passive action recognition^102^. The increased activation of the SMA and PMC during motor in imitation (vs. observation) suggests that these areas are consistent with well-established critical recognition of these regions being central to for internally generating motor sequences based on observed actions.

Furthermore, our findings reveal that there is a large degree of shared brain activation pattern between motor imitation and observation, particularly within the MNS (**Figure 3 and Supplementary Figure 3**). Both tasks recruit a shared network of brain regions involved in perception and motor planning, including the bilateral occipital lobe, STG, MTG, FFG, IFG and IPL (**Figure 3**). The concurrent activation of these areas suggests that the brain processes observed movements in a way that not only supports action understanding but also primes the motor system for potential execution. This functional overlap underscores the role of these regions in action encoding, reinforcing the idea that motor imitation and observation rely on a common neural substrate that bridges perception and movement.

While our findings align with previous literature on the neural basis of imitation as stated above, they also provide new insights into the task specific neural profiles during the gross motor observation and imitation as well as the evidence of a structured neural progression from posterior sensory regions to anterior motor areas ^103^. This highlights the hierarchical organization of motor learning, where initial sensory-driven processing in temporal-parietal regions is followed by predictive coding and motor planning in SMA and PMC ^99,104^. Additionally, the differential activation of the TPJ during observation versus imitation suggests additional neural recruitment from externally driven attention toward internally guided motor execution ^94,105^. These results suggest that imitation engages a specialized motor planning mechanism beyond passive observation, emphasizing the STG, SMA and PMC as critical hubs for transforming visual input into executable motor commands ^56,100,106^.

Beyond the MNS, our findings highlight the differential engagement of the VAN and dorsal attention network (DAN) across tasks. Our results reveal that there was greater VAN activation during observation than imitation (**Supplementary Figure 4, 5**), supporting its role in detecting and processing salient external stimuli ^74,75,77,89^. Additionally. the heightened activation of right TPJ (**Supplementary Figure 2**) suggests that passive observation of human actions relies heavily on stimulus-driven attention and social cue processing^94,95^. Conversely, the DAN (including IPS and frontal eye fields, FEF) was more engaged during imitation than observation, which aligns with its function in goal-directed attention and action planning (**Supplementary Figure 5**). The increased IPS activity during imitation might indicate that spatial and motor attention plays a crucial role in translating visual input into coordinated movements. This contrast between VAN and DAN activation aligns with previous findings that observation primarily involves passive stimulus-driven processing, whereas imitation requires active attentional control and motor planning.

The analysis of brain-behavior correlations between brain activation and CAMI scores revealed associations in MNS regions implicated in motor planning and execution. Positive correlations were observed in right parietal and left superior frontal gyrus, suggesting that greater motor imitation fidelity (i.e., higher CAMI scores) is linked to enhanced activation in these areas. Interestingly, we found the negative correlation in the right primary motor cortex and left middle temporal gyrus. This does not align with the hypothesis that stronger motor system engagement supports better imitation performance. However, the reduced activation in the right M1 could reflect greater efficiency in motor processing, where less activation may indicate that the brain is optimizing motor control without needing additional cortical resources^107,108^. An fMRI study found a positive correlation in the primary motor cortex with participants’ motor performance^109^. Another study found inferior parietal lobe (IPL; also implicated as part of the MNS) activity was negatively correlated with imitation accuracy^110^. Moreover, extant literature states that reduced activation in the core MNS regions may be associated with difficulties in motor adaptability^56,93,111^. These findings provide insight into how deficits in neural regions critical for motor control and imitation may contribute to reduced imitation fidelity in autistic individuals.

The relationship between brain activation and social responsiveness was further investigated for both motor observation and imitation. Positive correlations between SRS-2 t-scores and brain activation were observed in regions such as the right angular gyrus (r-AG) and left secondary visual cortex (l-SVC), particularly during observation. Although these correlations did not reach statistical significance during motor imitation, they provide preliminary evidence that impaired brain function in regions involved in action understanding and social cognition may underlie the social difficulties observed in ASD. Interestingly, our results show a positive correlation between SRS-2 and brain activation in right parietal lobe. This result diverges from our original hypothesis, which posited that brain response strength within the MNS during observation and imitation would be negatively correlated with dimensional measures of autistic traits. Instead of a reduced brain activation linked to higher autistic traits, our data indicate an enhanced reliance on the right parietal lobe in individuals with stronger autistic traits. This finding may indicate that individuals with higher autistic traits rely more heavily on this region to process and understand observed actions or social cues, potentially compensating for difficulties in other aspects of social interaction^112^..

### Limitations

While this study provides valuable insights into neural correlates of motor observation and imitation and its association with autistic traits, several limitations should be addressed in future research. First, the relatively small sample size of the ASD group limits the generalizability of the findings in autistic individuals. Rather, we used a dimensional approach to analysis and incorporated ASD samples that allowed us to represent a broad range behavioral measurements and conduct the brain behavioral association under of motor observation imitation Future studies with larger and more diverse populations are needed to confirm and extend these results. Second, although HD-DOT offers significant advantages over traditional neuroimaging methods, its field of view is less extensive than that of fMRI, which limits sensitivity to neural activity in deeper regions such as cerebellum and sub-cortical regions previously implicated in ASD functional neuroimaging studies^113,114^. Third, we did not use eye tracking to monitor eye-gaze and visual attention throughout the experiment. Instead, participants were instructed to focus their attention on the screen, and they were continuously visually monitored for attention and imitation performance by the lab member conducting the study. Such simultaneous collection of eye gaze with neuroimaging and 3D motion tracking will provide a powerful strategy for future studies to more fully investigating the role of visual attention on motor output. Fourth, in this study, data loss from the initial 100 participants occurred due to independent exclusions across different data types, including HD-DOT quality, missing SRS-2 scores, and CAMI recording issues. These exclusions did not systematically affect the same participants across modalities or groups. Finally, although the current study focused on adults, future studies that focus on how brain function supports naturalistic motor imitation throughout childhood development, including in children with and without autism, will be positioned to inform brain-region-targeted interventions and advance precision medicine. Those investigation will provide valuable insights into the developmental aspects of naturalistic motor imitation, shedding light on how these mechanisms may differ throughout the lifespan and how they may contribute to the social and motor challenges observed in children with ASD.

## Conclusion

In this study, we used simultaneous HD-DOT functional neuroimaging with the CAMI method to investigate how brain function covaries with naturalistic motor observation and imitation. Consistent with our hypotheses based on prior fMRI studies under less naturalistic conditions, our findings reveal commonality of neural activation patterns for motor observation and imitation, principally within MNS regions. We also observed some clear distinctions in neural activation patterns between passive action observation and active motor imitation. Observation relies more on temporal-parietal regions (e.g., STG, IPL, TPJ) and the VAN, reflecting stimulus-driven processing of movement cues. In contrast, imitation engages prefrontal and motor regions (M1, SMA, PMC, IFG) and the DAN, highlighting the additional demands of motor planning and execution. The additional neural recruitment from posterior (sensory-driven) to anterior (motor-driven) cortical activation during imitation underscores the neural transformation from perception to action execution. Moreover, our brain-behavior analyses indicated that reduced MNS activation may associate with higher autism-related traits and lower motor imitation fidelity. Overall, these findings highlight the neural correlates of action observation/execution, associated social reciprocity and motor imitation fidelity. Finally, these studies lay essential foundation for future investigations using simultaneous methods for naturalistic functional neuroimaging and quantitative CAMI that may be applied during early childhood screenings for ASD.,

## Acknowledgements

We would like to express our sincere gratitude to the participants for their valuable contributions to this study. This research was supported by grants from the National Institutes of Health (R01-MH12275101, R21-MH127501), the McDonnell Center for Systems Neuroscience, the Simons Foundation for Autism Research (SFARI) and the Eagles Autism Foundation. All study procedures were reviewed and approved by the Human Protections Office at the Washington University School of Medicine Institutional Review Boards. Written informed consent was obtained from all participants. Additionally, the individuals showed in the image in Fig. 1 provided consent for the use of their photo in publications.

## Data and Code Availability

The datasets generated and analyzed during the current study, as well as the analysis codes, are available from the corresponding author upon request. All fully de-identified data will also be available on the NIMH data archive (NDA).

## Author Contributions

Conceptualization: ATE, SHM. Data collection: DY, TGG, CMS, SRM, KTK, EDD, SP, ES, AS. Formal analyses: DY, TGG, ATE. Methodology: DY, CP, RR, DL, RS, DC, ADS, MBN, BT, RV, NM, SHM, ATE. Project administration: ATE, SHM. Resources: ATE, SHM. Software: ATE, SHM, BT. Visualization: DY, ATE. Writing: DY, ATE. Funding acquisition: ATE, SHM. All authors read and approved the final manuscript.

---

**Supplementary Figure 1:**
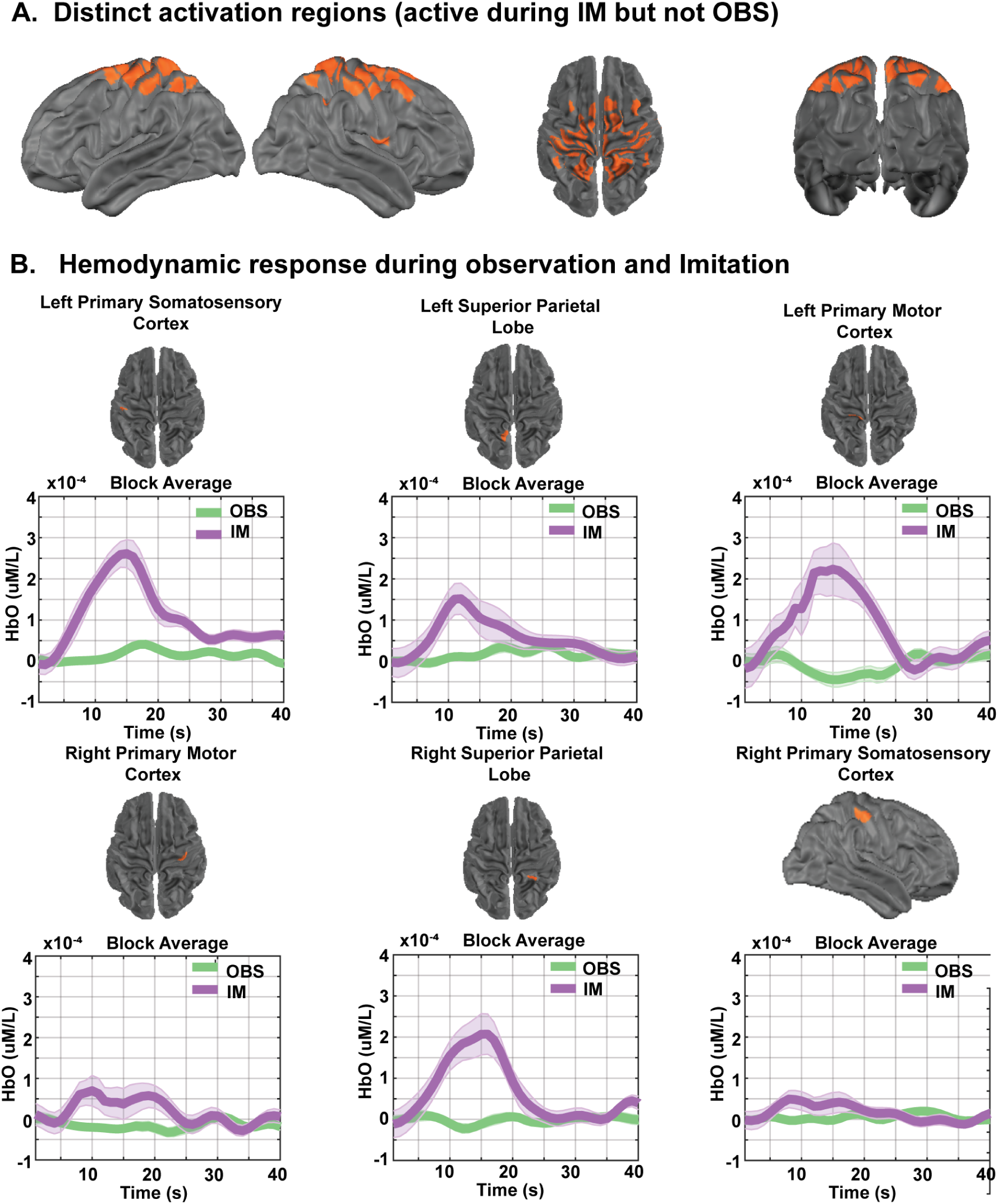
Brain regions exhibiting significant responses during motor imitation but not motor observation. **A.** Brain regions with distinct significant activation (FDR corrected *p* < 0.05) during motor imitation that are not significantly active during motor observation. **B.** Temporal profiles of the HbO (oxygenated hemoglobin) hemodynamic response for each task, with shaded areas representing the standard error of the mean across participants.

**Supplementary Figure 2:**
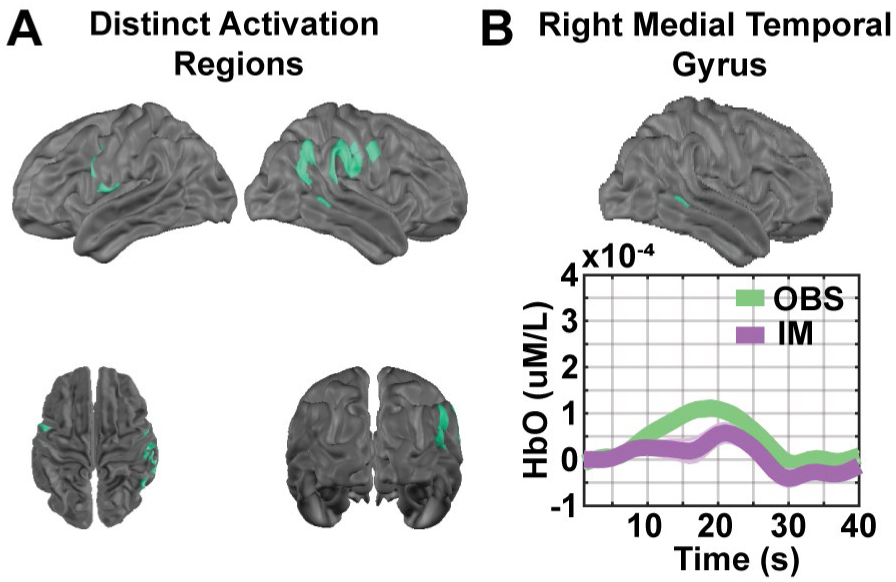
Brain regions exhibiting significant responses during motor observation but not motor imitation. **A.** Brain regions with distinct significant activation (FDR corrected *p* < 0.05) during motor observation task that are not significantly active during motor imitation. **B.** Temporal profiles of the HbO (oxygenated hemoglobin) hemodynamic response for each task, with shaded areas representing the standard error of the mean across participants

**Supplementary Figure 3:**
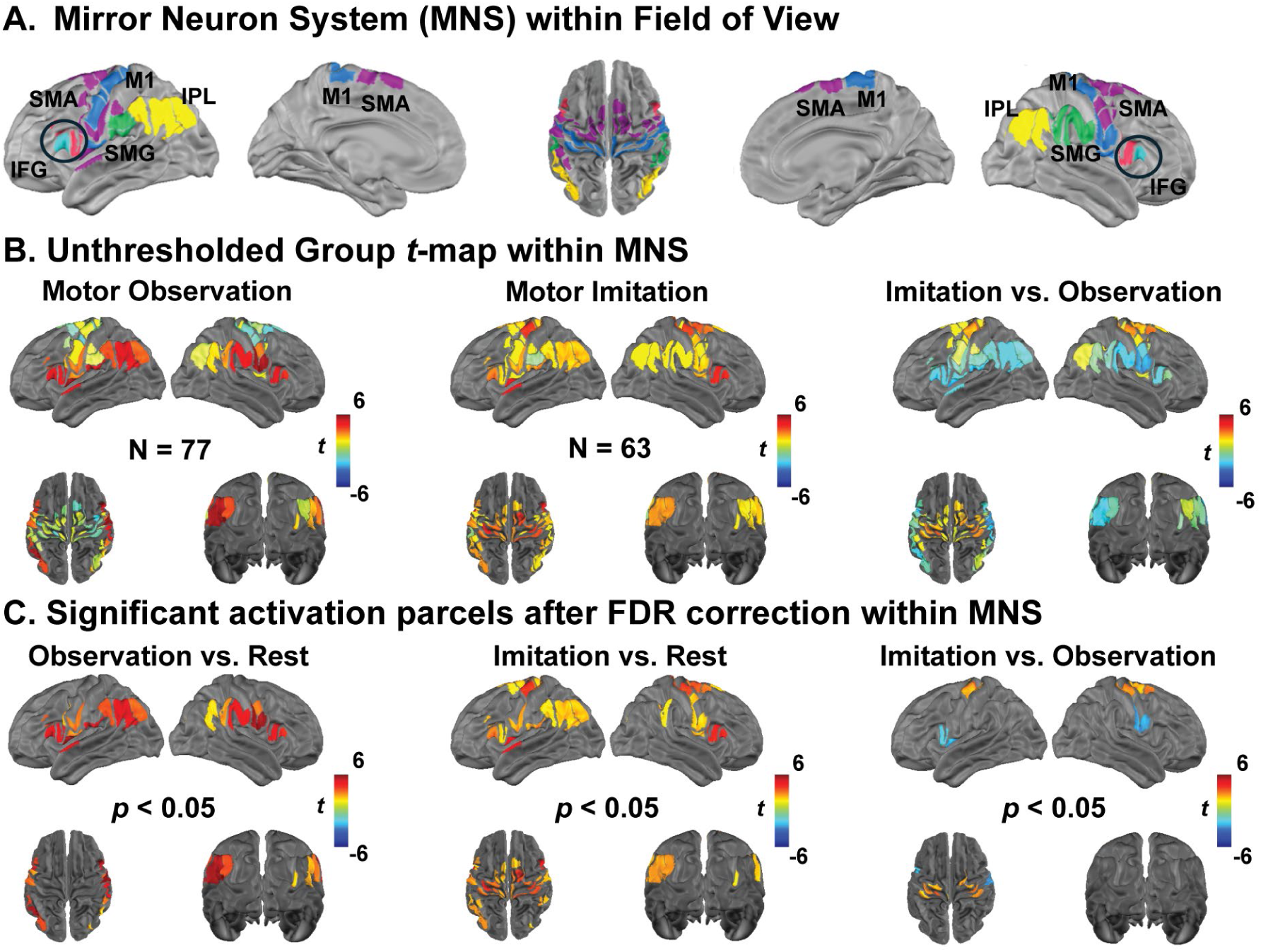
Neural correlates of observation and imitation within the mirror neuron system. A. Parcel-based analysis of the MNS within the field of view, highlighting regions activated during both motor observation and imitation. B. FDR-corrected t-map showing the statistically significant regions within the MNS, with false discovery rate correction applied to control for multiple comparisons.

**Supplementary Figure 4:**
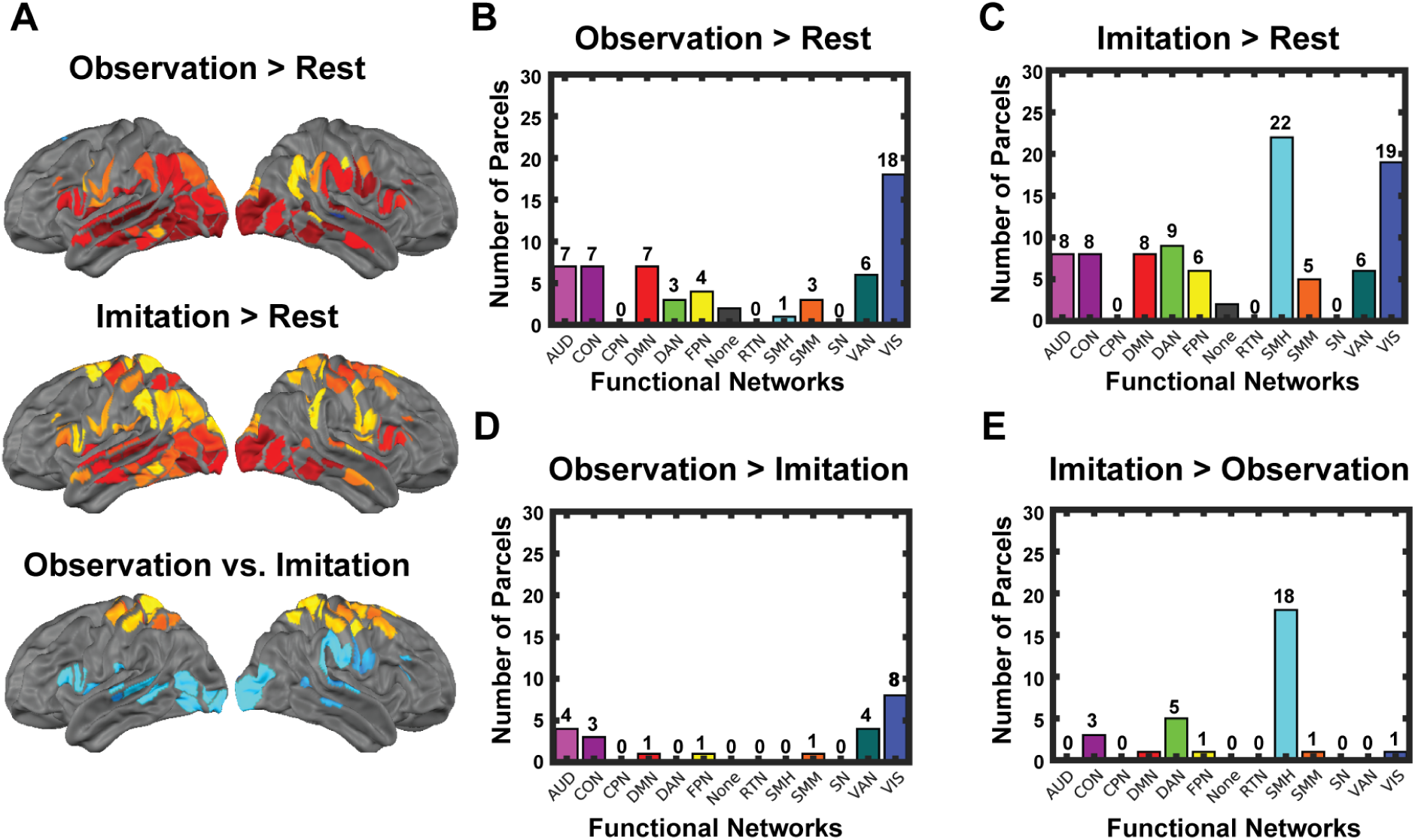
Functional network involvement during motor observation and imitation. **A** Task specific significant brain parcels under three conditions (OBS, IM, and their contrast). **B** Functional networks involved during the OBS task. **C** Functional networks involved during the IM task. **D** Functional networks more active during the OBS task than the IM task. **E** Functional networks more active during the IM task than the OBS task.

**Supplementary Figure 5:**
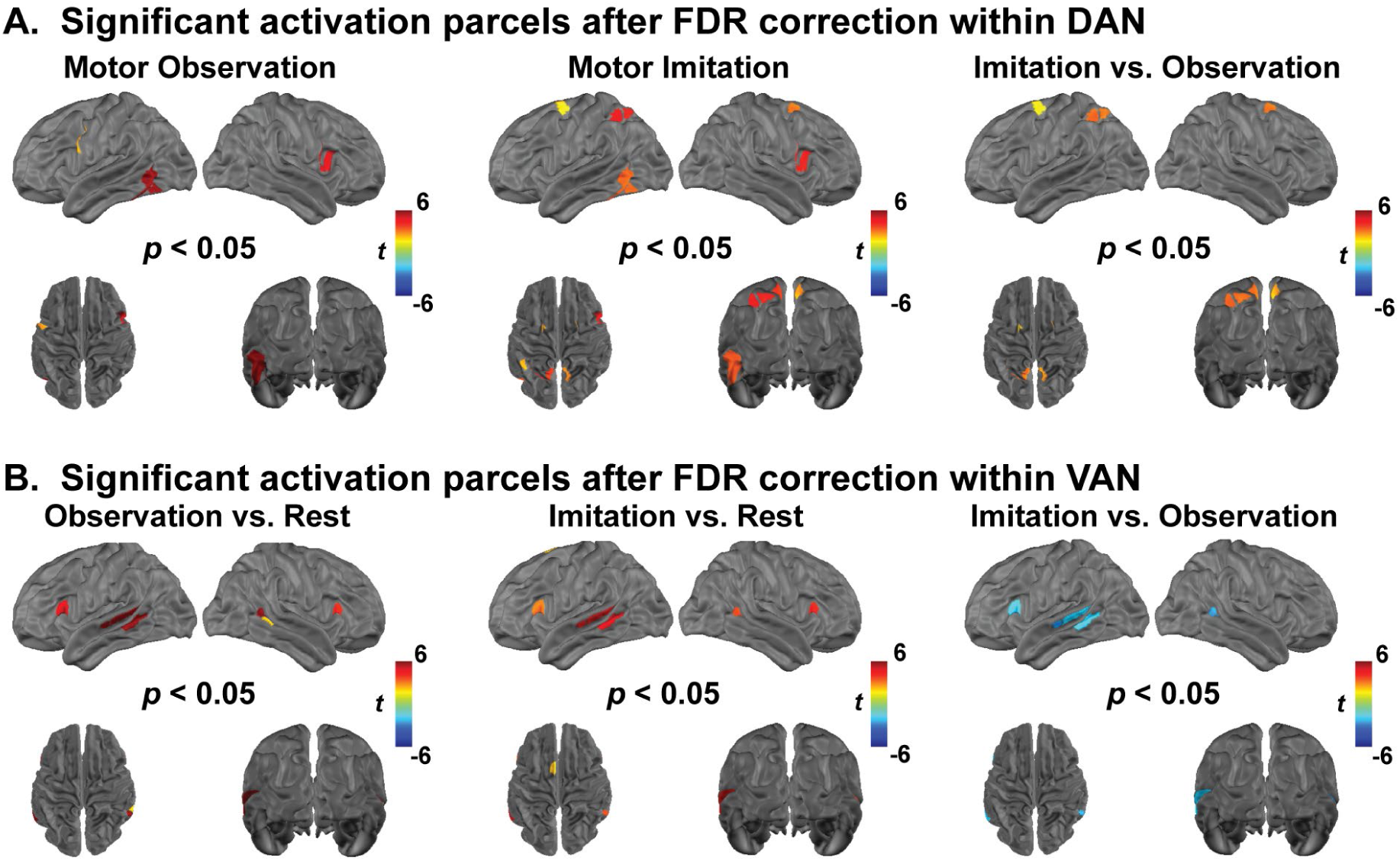
Neural correlates of OBS and IM within dorsal attention network (DAN) and ventral attention network (VAN). **A.** FDR-corrected t-map for motor observation (N = 77), motor imitation (N = 63), and their contrast (red indicates brain regions where OBS > IM, highlighting statistically significant regions within DAN with false discovery rate correction applied to account for multiple comparisons. **B.** FDR-corrected t-map for motor observation (N = 77), motor imitation (N = 63), and their contrast (red indicates brain regions where OBS > IM, highlighting statistically significant regions within VAN with false discovery rate correction applied to account for multiple comparisons.

